# Recovering genomes and phenotypes using allele-specific gene expression

**DOI:** 10.1101/2020.11.11.377978

**Authors:** Gamze Gürsoy, Nancy Lu, Sarah Wagner, Mark Gerstein

## Abstract

With the recent increase in RNA sequencing efforts using large cohorts of individuals, studying allele-specific gene expression is becoming increasingly important. Here, we report that, despite not containing explicit variant information, a list of allele-specific gene names of an individual is enough to recover key variants and link the individual back to their genome or phenotype. This creates a privacy conundrum.

---

Owing to the surge in functional genomics data over the past decade, numerous reports have focused on identifying genomic privacy issues related to molecular endophenotype data, such as gene expression levels^1,2^. These studies exploit the known and publicly available relationship between genotype and endophenotypes such as expression quantitative trait loci (eQTLs). That is, given a matrix of gene expression values collected from a cohort of individuals and a list of eQTLs, one can link a genome from a known individual to the gene expression matrix and uncover potentially stigmatizing phenotypes such as HIV status^1^.

The increase in phased personal genomes and functional genomics data allows researchers to investigate the allele-specific activity of the genome. With the surge in large-scale RNA sequencing and genotype efforts such as the Genotype-Tissues Expression (GTEx) project^3,4^, more studies have begun focusing on allele-specific expression (ASE) in the human genome^4–7^. ASE is a characteristic of expressing only one copy of a gene (maternal or paternal allele) and may lead to phenotypic variation. Between 10 and 22% of human genes show allele-specific regulation of gene expression^8^. ASE can be created in part by underlying biological processes such as imprinting. However, most observed cases are not necessarily due to an underlying biological phenomena. There is increasing evidence that ASE could be linked to the predisposition to diseases such as autism spectrum disorder^9^, colorectal cancer^10^ and, tumorigenesis in general^11^.

Due to their clinical importance and direct relationship to the phenotype of the organism, there is an incentive to broadly share a list of allele-specific genes, or allele-specific gene expression matrices, of study participants. Moreover, ASE information is often shared with the accompanying phenotype. Many assume that haplotype-level gene expression data do not contain any identifying information and are safe to share even if the data are derived from individuals who did not provide broad consent^4^.

Here, we demonstrate that privacy breaches are possible solely by using a list of allele-specific gene names of an individual. As an example, we show these breaches using ASE data from individuals of the 1,000 Genomes Project, in which the full genomes of the individuals are broadly shared.

Genomic privacy attacks in the form of linking two datasets together can be categorized differently based on whether the genomic variants observed from the linked datasets are noisy or perfect^12^. However, the privacy attacks we describe here differ in nature from previous linkage attacks^1,2,12,13^, as the genomic variants cannot directly be observed from the ASE data (Figure S1). Two privacy attacks can be performed using an ASE gene name list: (1) Recovering the genome of an individual; and (2) inferring the phenotype of an individual.

In the first attack, the adversary obtains a list of ASE gene names of a known individual (perhaps through electronic health records or a friendly conversation). The goal is to link these gene names to an anonymized publicly available genome dataset to recover the genome of the known individual (Figure 1A).

**Figure 1:**
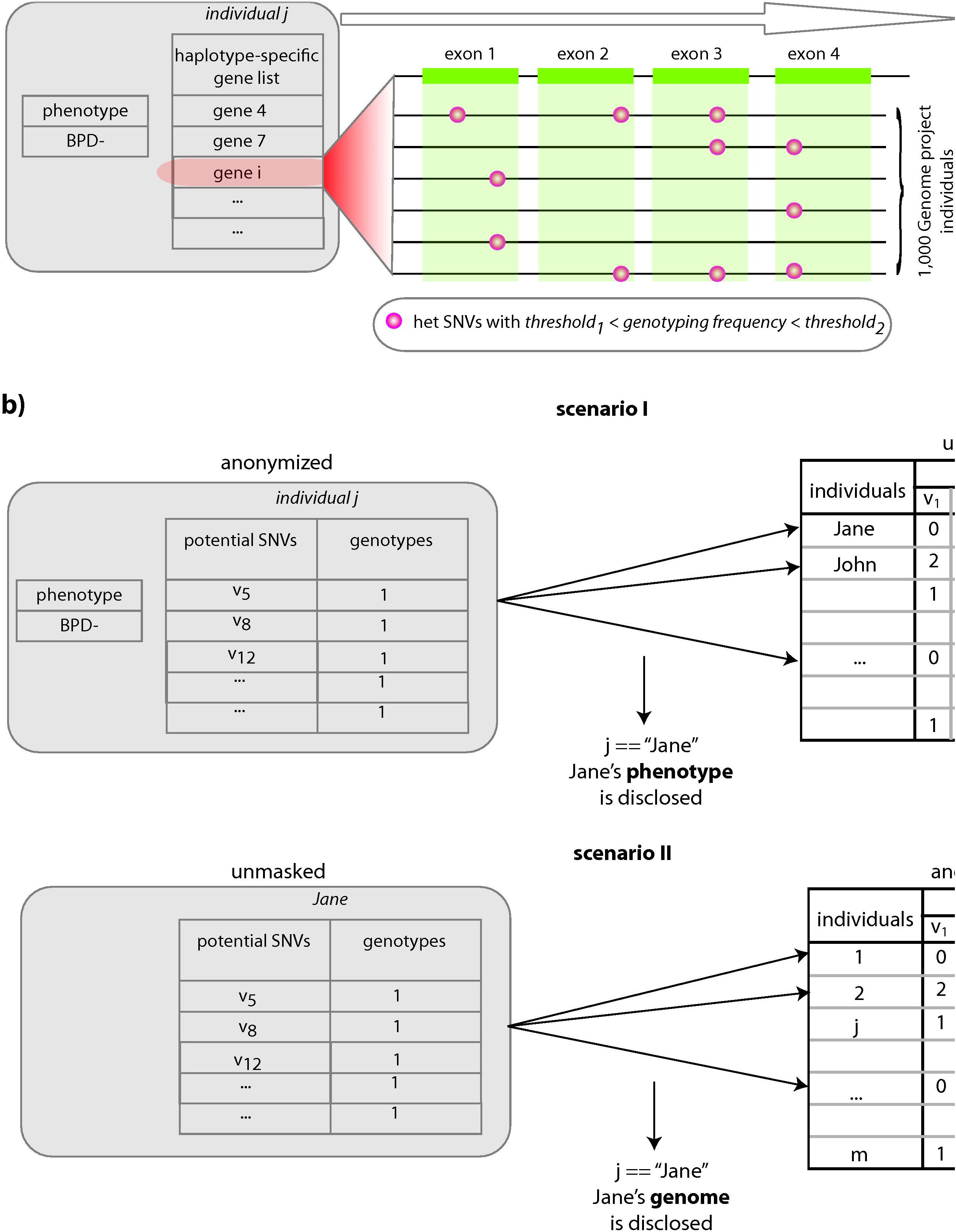
Schematic representation of using allele-specific genes to de-anonymize individuals. a) Schematic of going from a list of genes to a list of SNPs. b) De-anonymizing a list of anonymous ASE genes using publicly available genomes from known individuals and inferring private phenotypes. b) Recovering the anonymized genome of a known individual by using their ASE gene list.

In the second attack, a research study (e.g., GTEx, PsychENCODE) releases a publicly available anonymized database. This database contains the allele-specific expression of all genes for a number of individuals. It also contains the sensitive phenotypes of these individuals. The adversary compiles a database of genomes from known individuals (e.g., those who participated in a research study), by using genetic genealogy databases such as GEDmatch. The goal of the adversary is to uncover the phenotypes of these known individuals of interest. The allele-specific gene expression database can be summarized as a list of ASE gene names for each individual in the database by comparing the gene expression between alleles. The adversary uses the genomes to mine the ASE gene name lists and finally links the genomes of known individuals to the phenotypes of the anonymized individuals (Figure 1B).

As mentioned before, there is no explicit genotype information in a gene name. However, if a gene is determined to be allele-specific for an individual, then an accessible heterozygous single-nucleotide polymorphism (SNP) must be present somewhere on the gene body such that researchers were able to phase the gene expression into alleles. By using this information, we overlapped the exon locations of the reported ASE genes with heterozygous SNPs in a database of genomes. This allowed us to generate a candidate SNP/genotype list for each ASE gene (Figure 1A, see Online Methods for details). Next, we used a linking approach^12,14^ that weights the SNPs according to their frequency in the database. This approach scored every individual in the database based on the similarity between the genome and the candidate genotypes either by using a best matching linking score approach^12^ or by using a probability distribution through entropy calculations^14^.

We used lists of allele-specific gene names from 382 individuals^6^ and attempted to link them to a database of 2,504 individuals^15^. We were able to link 55% of the individuals to the database using only a list of allele-specific gene names per individual (Figure 2a). This number increased to 80% when we relaxed our criteria from the best-matching individual to any individual in the top 20 best matches (Figure 2b).

**Figure 2.**
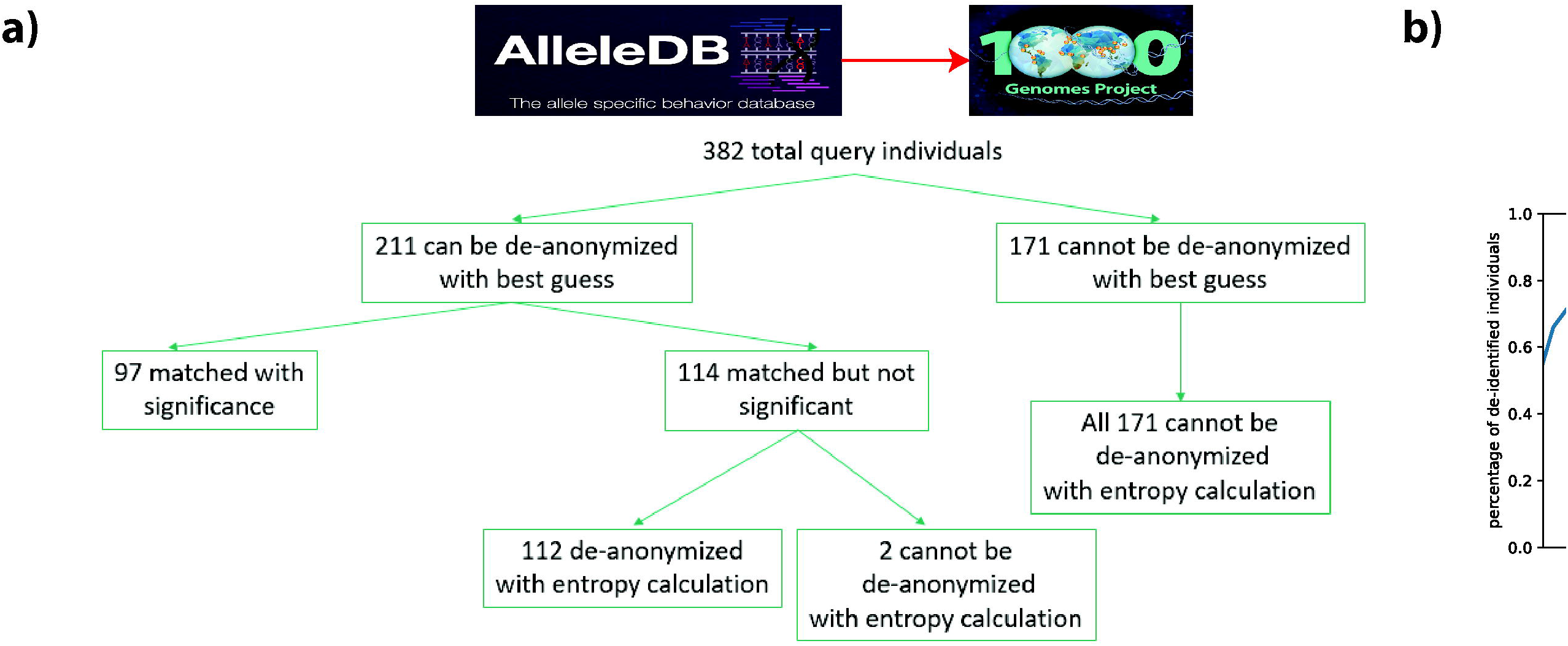
Linking attack accuracy. a) The number of individuals that can be linked to their genomes with different statistical techniques. b) The percentage of individuals that can be linked to their genomes when we relax the criteria from best match to top k ranked.

Next, we found that highly polymorphic human leukocyte antigen (HLA) genes were in the majority of the individuals’ gene lists (Figure 3a). We first used only HLA genes for the linking and found that we could link only 2.9% of the individuals to the database. We then removed the HLA genes from the original ASE gene list of each individual and found that the percentage of correctly linked individuals increased from 55% to 66%. Lastly, we removed the top 20 genes that were common to the individuals in the database from the gene list. This further increased the percentage of linked individuals to 68% (Figure 3b).

**Figure 3.**
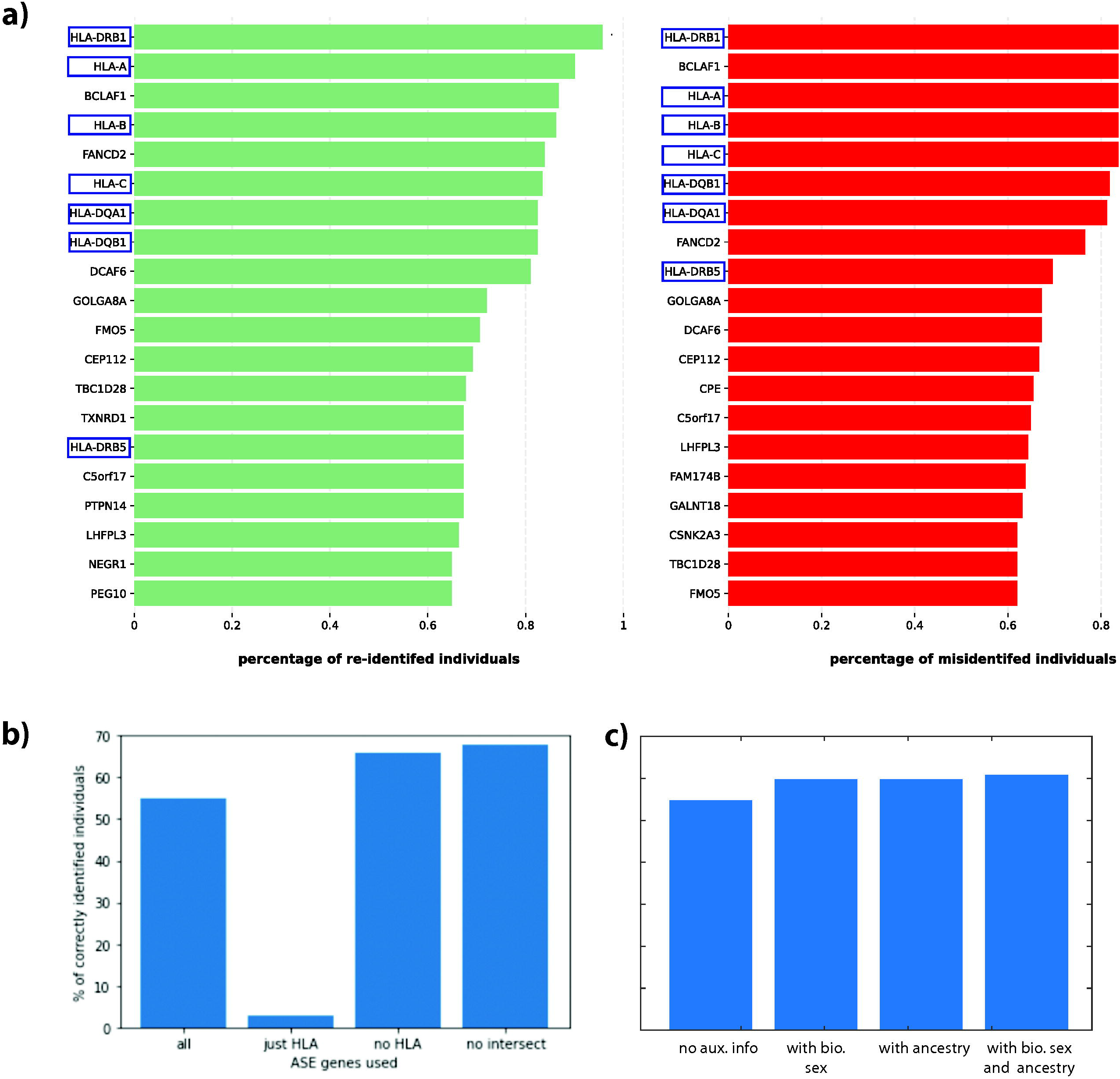
Impact of auxiliary information on linking ability. a) The top 20 genes that are found on the ASE gene list of correctly identified and misidentified individuals. b) The percentage of correctly linked individuals when we used different combinations of ASE genes. c) The percentage of correctly linked individuals when we used biological sex and/or ancestry as auxilary information.

Our last test was to calculate the improvement in linking when we used auxiliary data. Knowledge of the biological sex of the individuals increased the percentage of correctly linked individuals to 60%. Adding only the ancestry information of the study participants also increased the percentage of correctly linked individuals to 60%. Although, knowledge of both ancestry and biological sex increased the percentage of correctly linked individuals by only ~1% (Figure 3c), the percentage of correctly linked individuals with high statistical significance increased by ~10% (Figure S2). We did not find a significant difference in the number of genes used per individual in the correctly linked and mislinked categories; however, when we weighted the number of genes by their length, we found that correctly linked individuals had longer ASE genes (Figure S3 a,b). We also did not find a significant difference in the number of candidate SNPs per individual between the correctly linked and mislinked categories (Figure S3 c).

This study shows that although ASE does not explicitly reveal the location of the SNPs of an individual, by using simple and straightforward biological knowledge can enable ASE genes to be linked to the genomes and/or phenotypes of study individuals. We showed the feasibility of this breach with data from individuals who provided broad consent. However, we envision that the same publicly available data could be used to infer private genetic variants of individuals who do not wish to release their genomes broadly. Furthermore, these inferred SNPs can lead to imputation of other genetic markers through linkage disequilibrium, which, in turn, might lead to even bigger privacy issues.

As is the case with other molecular phenotype and functional genomics data, preventing the public release of ASE genes can hamper biomedical discoveries and clinical studies. Researchers could perform risk assessments of releasing gene names based on their polymorphism and length. Based on this assessment, some genes could be omitted from the list. However, this approach might reduce the utility of the released data. We believe that the best approach to mediate genomic privacy issues related to hidden information in summary-level functional genomics data is three-pronged: (1) develop clear and detailed informed consent policies, (2) educate participants on the risks and benefits of the study, and (3) establish laws and legislation to prevent bad actors from using genetic information to harm individuals, as noted by earlier genomic privacy studies^16^.

## Data and software availability

The code used in this study can be found at https://github.com/gersteinlab/privaseq4. 1000 Genomes vcf files are downloaded from ftp sites in https://www.internationalgenome.org. The processing steps and example files can be found in the data folder of the github page. A gene list for each individual is taken from http://alleledb.gersteinlab.org and can be found in the data folder of the github page. The locations of the exons for these genes are also provided in the data folder of the github page.

## Methods

### Compiling a list of candidate SNPs from the ASE gene list

We first overlapped the gene names with the genes in the GENCODE comprehensive gene annotation file (release 19, GRCh37.p13) to pinpoint the location of the exons of these genes. For each gene, we found all the SNPs that overlapped with its exons by using a vcf file from 2,504 individuals (1,000 Genomes database). We first calculated the heterozygous genotyping frequencyof these SNPs as 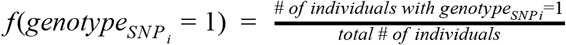. We then removed the SNPs that had 0.1< *f*(*genotype_SNP_i__* = 1) < 0.5 from the overlap list. We added the remaining SNPs to the candidate SNP list. We repeated this procedure for all of the genes in the list to obtain one final candidate SNP list.

### Linking attacks

Let us assume that we have *n* total SNPs that can be observed in humans (e.g., all of the SNPs observed in the 1,000 Genomes Project). We can represent an individual’s genome as a set *S* = {*g*_1_, *g*_2_,…, *g_n_*}, where *g_i_* is the genotype of the *i^th^* SNP. Candidate SNPs obtained using ASE genes become a subset of S, whose genotypes are assumed to be heterozygous (*g_i_* = 1), i.e., *S_can_* = {*g*_1_ = *x, g*_2_ = 1,…, *g_n_* = *x*}, where *g_i_* = *x* means SNP *i* is not in the candidate list, hence its genotype is unknown.

### Scenario 1

Let us assume we have an ASE gene list of a known individual. This means we can compile a list of heterozygous SNPs for this known individual. In this case, *S_can_* = {*g*_1_ = *x, g*_2_ = 1,…, *g_n_* = *x*} is the set of candidate genotypes for the known individual. The goal is to recover the genotypes for all of the SNPs in the set. Let us assume we have access to a database of anonymized genomes. Each anonymized genome *j* in the database can be represented as 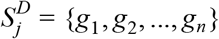, where each genotype *g_i_* is known.

#### Best match approach

For each individual *j* in the database, we find the intersection 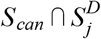 and calculate a linking score 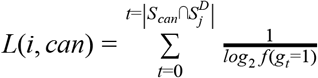, where *f*(*g_t_* = 1) is the ratio of individuals whose *t^th^* SNP has the heterozygous genotype (*g_t_* =1) to the total number of individuals in *D* [previously defined in^12^]. To recover the genome for the known individual, we then rank all the *L*(*i, can*) scores for all genomes in *D* in decreasing order. We denote the genome with the highest score as the genome of the known individual with candidate SNPs. To assess the statistical robustness of this prediction, we used our previously defined *gap* measure, which is the ratio between the highest and second highest *L*(*i, can*) scores. We further calculate the statistical significance of *gap* by generating random candidate SNPs (as many as the original candidate SNPs), perform the above attack one thousand times, and compare the real *gap* value against the distribution of random *gap* values.

#### Entropy approach

The goal of this approach is to assign a probability of correctly linking the ASE list to each genome in *D*, which allows us to have a distribution. This approach is adopted from Narayanan and Schamtikov, 2012. We calculate the probability of linking the candidate SNP list to a genome *i* in *D* as 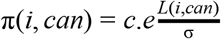, where *c* is a constant to satisfy 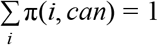, *L*(*i, can*) is the linking score described above, and *σ* is the standard deviation of the linking scores (Figure S4).

### Scenario 2

The mathematical formulation of scenario 2 is the same as the first scenario. The only difference is that we have the genome of the known individual and we try to link this known genome to an anonymized ASE gene list, which is connected to a potentially private phenotype.

### Identification of the top 20 common genes

After linking 382 ASE gene lists to a genome in *D*, we calculated the accuracy of the linking. We then separated the gene lists into two categories: (1) lists that led to correct re-identification and (2) lists that led to misidentification. We identified the genes that were shared across many ASE gene lists in both categories. Among the top 20 shared genes, we found that HLA genes were in the lists of >90% of both correctly re-identified and misidentified individuals. We then selectively removed different groups of genes (HLA, and genes at the intersection of both groups) and performed the linking attacks.

### Usage of auxiliary data

We added one or two more features to our sets 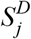 (the genotypes of genome j in database D) and *S_can_* (the candidate SNP genotype list) such that our new list does not only have genotypes but also includes biological sex and/or ancestry features. 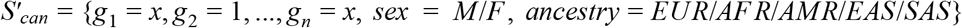 and 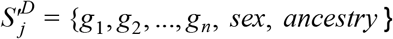 are our new sets and we look for the 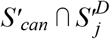 intersection to calculate the linking scores. Here, M and F are used for biologically male and female individuals, respectively. EUR, AFR, AMR, EAS, and SAS correspond to European, African, Admixed American, East Asian, and South Asian ancestries, respectively.

## Notes

### Competing Interest Statement

The authors have declared no competing interest.

