## Supplemental Figures for "Recovering genomes and phenotypes using allele-specific gene expression"

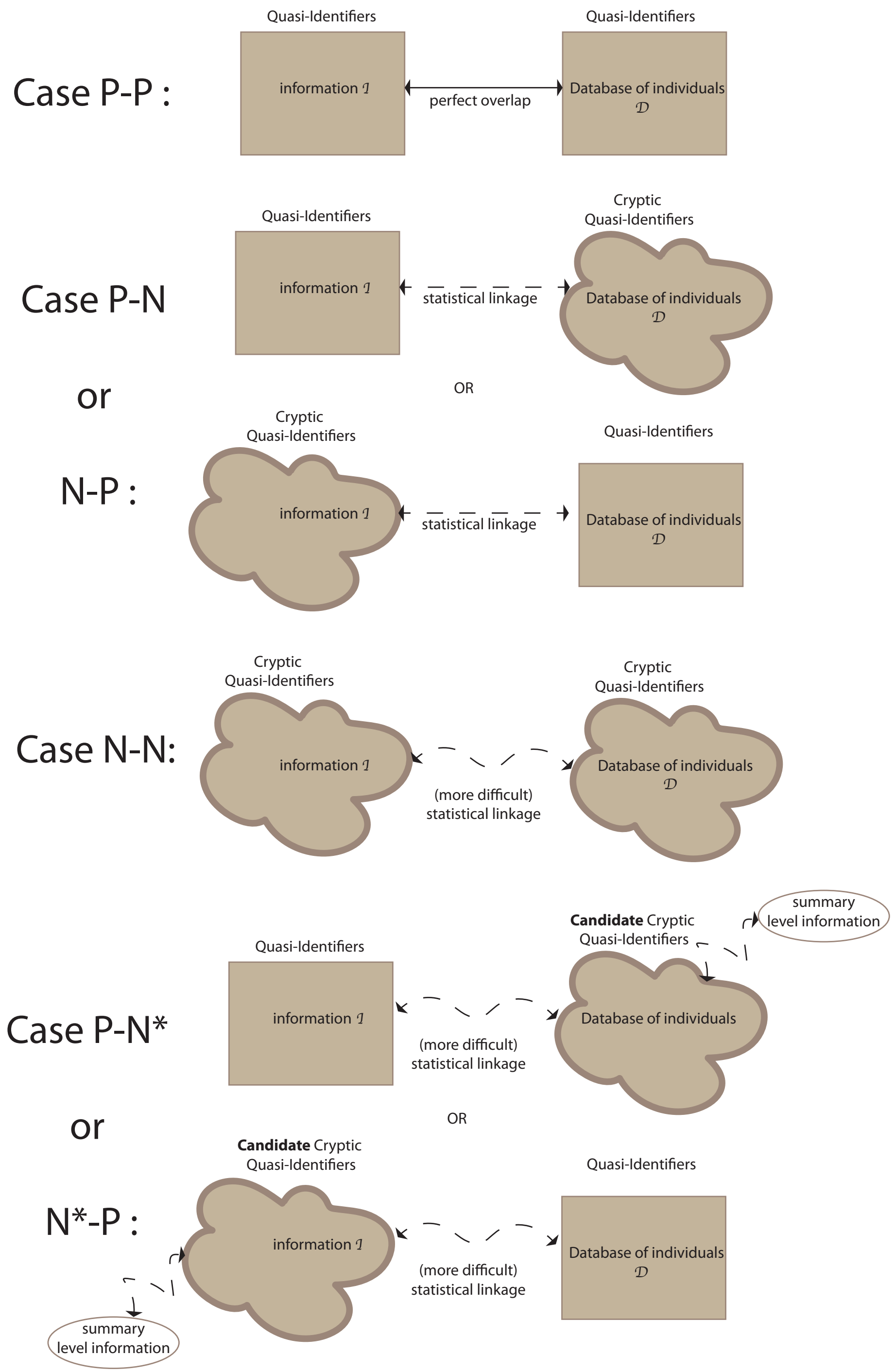


Figure S1: Different cases of linkage attacks

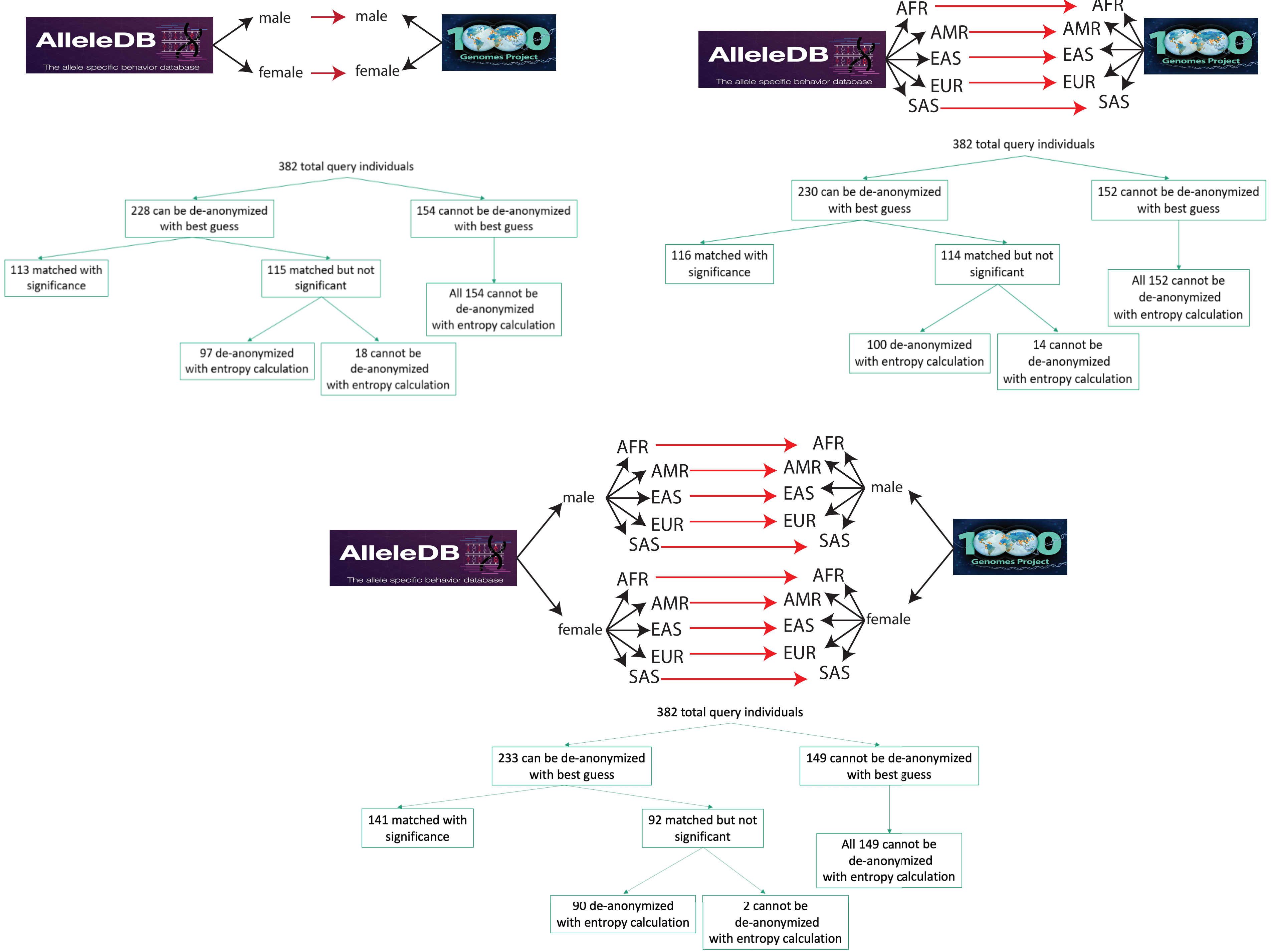


Figure S2: Number of individuals that are correctly identified using different methodologies and different auxiliary information.

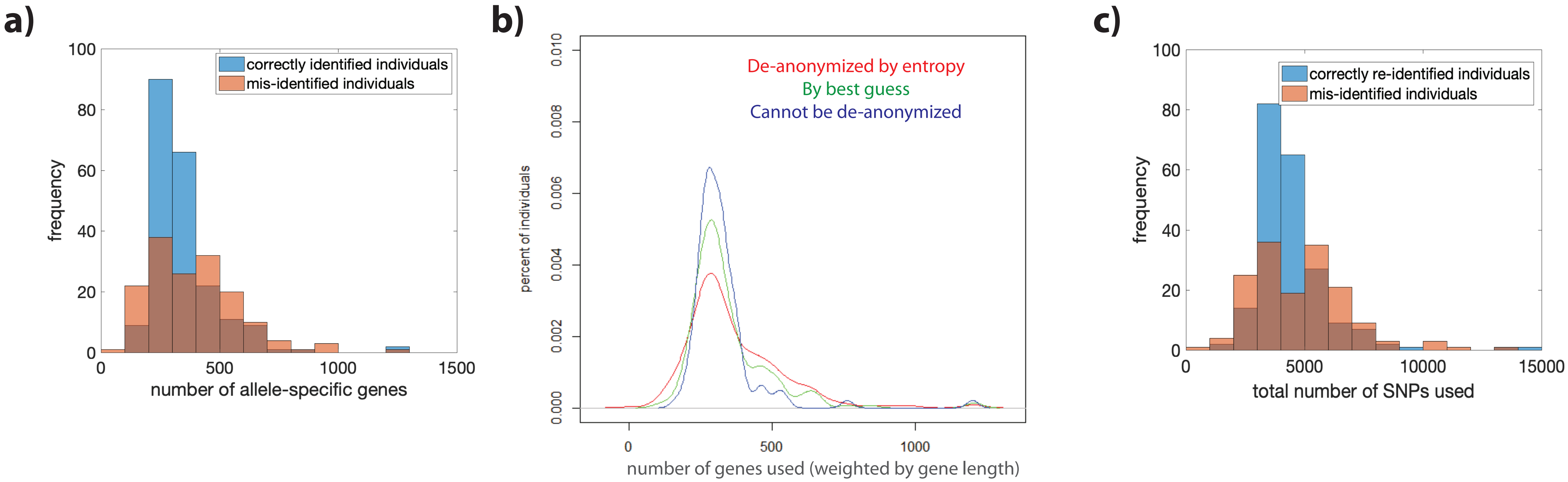


Figure S3: **a)** Distribution of number of ASE genes used for correctly identified individuals (in blue) and for mis-identified individuals (in orange). **b)** Same as (a), but the genes are weighted by their length. **c)** Distribution of number of candidate SNPs inferred for correctly identified individuals (in blue) and for mis-identified individuals (in orange).

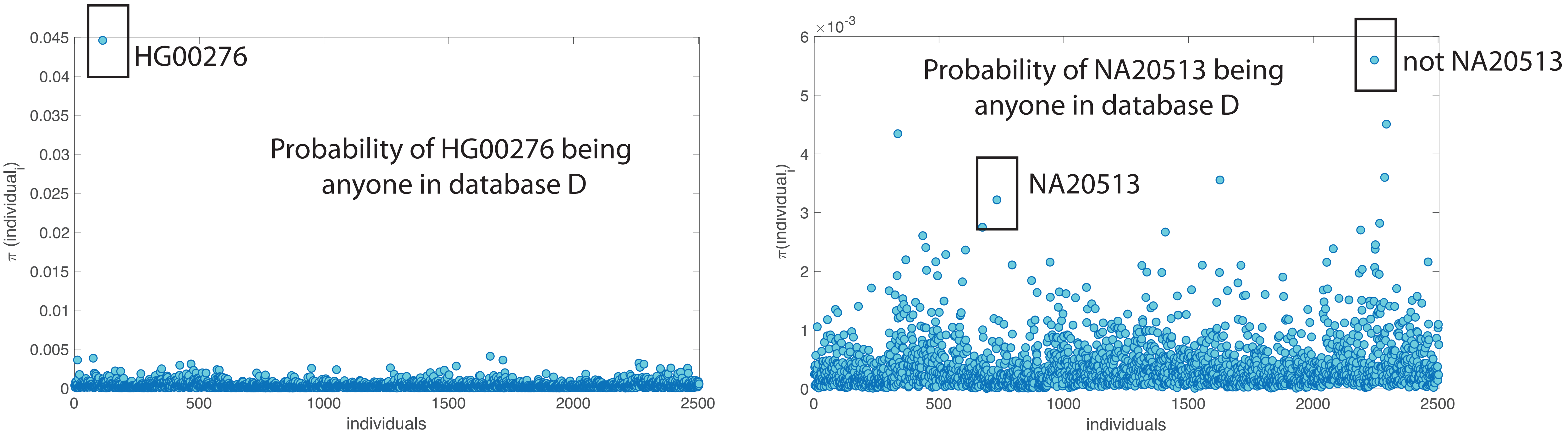


Figure S4: Examples of entropy based matching for a correctly identified individual (HG00276) and mis-identified individual (NA20513)
